# Rational Thoughts in Neural Codes

**DOI:** 10.1101/765867

**Authors:** Zhengwei Wu, Minhae Kwon, Saurabh Daptardar, Paul Schrater, Xaq Pitkow

## Abstract

Complex behaviors are often driven by an internal model, which integrates sensory information over time and facilitates long-term planning to reach subjective goals. We interpret behavioral data by assuming an agent behaves rationally — that is, they take actions that optimize their subjective reward according to their understanding of the task and its relevant causal variables. We apply a new method, Inverse Rational Control (IRC), to learn an agent’s internal model and reward function by maximizing the likelihood of its measured sensory observations and actions. This thereby extracts rational and interpretable thoughts of the agent from its behavior. We also provide a framework for interpreting encoding, recoding and decoding of neural data in light of this rational model for behavior. When applied to behavioral and neural data from simulated agents performing suboptimally on a naturalistic foraging task, this method successfully recovers their internal model and reward function, as well as the computational dynamics within the neural manifold that represents the task. This work lays a foundation for discovering how the brain represents and computes with dynamic beliefs.

---

Understanding how the brain works requires interpreting neural activity. The behaviorist tradition (1) aims to understand the brain as a black box solely from its inputs and outputs. Modern neuroscience has been able to gain major insights by looking inside the black box, but still largely relates measurements of neural activity to the brain’s inputs and outputs. While this is the basis of both sensory neuroscience and motor neuroscience, most neural activity supports computations and cognitive functions that are left unexplained — we might call these functions ‘thoughts’. To understand brain computations, we should relate neural activity to thoughts. The trouble is, how do you measure a thought?

Here we propose to model thoughts as dynamic beliefs that we impute to an animal, by combining explainable Artificial Intelligence (AI) cognitive models for naturalistic tasks with measurements of the animal’s sensory inputs and behavioral outputs. We define an animal’s task by the relevant dynamics of its world, observations it can make, actions it can take, and the goals it aims to achieve. The AI models that solve these tasks generate beliefs, their dynamics, and actions that reflect the essential computations needed to solve the task and generate behavior like the animal. With these estimated thoughts in hand, we propose an analysis of brain activity to find neural representations and transformations that potentially implement these thoughts.

Our approach combines the flexibility of complex neural network models while maintaining the interpretability of cognitive models. It goes beyond black-box neural network models that solve one particular task and find representational similarity with the brain (2–4). Instead, we solve a whole family of tasks, and then find the task whose solution best describes an animal’s behavior. We then associate properties of this best-matched task with the animal’s mental model of the world, and call it ‘rational’ since it is the right thing to do under this internal model of the world. Our method explains behavior and neural activity based on underlying latent variable dynamics, but it improves upon usual latent variable methods for neural activity that just compress data without regard to tasks or computation (5–7). In contrast, our latent variables inherit meaning from the task itself, and from the animal’s beliefs according to its internal model. This provides interpretability to both our behavioral and neural models.

We also want to ensure we can explain crucial neural computations that underlie ecological behavior in natural tasks. We can accomplish this by using tasks with key properties that ensure our model solutions implement these neural computations. First, a natural task should include latent or hidden variables: animals do not act directly upon their sensory data, as that data is merely an indirect observation of a hidden real world (8). Second, the task should involve uncertainty, since real-world sense data are fundamentally ambiguous and behavior improves when weighing evidence according to its reliability. Third, relationships between latent variables and sensory evidence should be nonlinear in the task, since if linear computation were sufficient then animals would not need a brain: they could just wire sensors to muscles and compute the same result in one step. Fourth, the task should have relevant temporal dynamics, since actions affect the future; animals must account for this.

While natural tasks that animals perform every day do indeed have these properties, most neuroscience studies isolate a subset of them for simplicity, such as two-alternative forced choice tasks, multi-armed bandits, or object classification. Although these have revealed important aspects of neural computation, they also miss some of the fundamental structure of brain computation. Recent progress warrants increasing the naturalism and complexity of the tasks and models.

One major challenge for practical studies with increased complexity and naturalism is to record from many neurons with enough spatial and temporal precision to reveal the relevant computational dynamics for these tasks. Specifically, the dimensionality of neural data needs to be bigger than the dimensionality of our target tasks (9). Modern neurotechnology now affords us this opportunity: brain-wide calcium imaging at cellular resolution and fine-grained electrophysiological recording can record from thousands of neurons simultaneously at high frequency. Limited experimental time and coverage still hinder our ability to explore the neural representations. But with current large-scale neural data, we will increasingly have enough power to find neural representations and dynamics in naturalistic and cognitively interesting tasks.

This paper makes progress towards understanding how the brain produces complex behavior by providing methods to estimate thoughts and interpret neural activity. We first describe a model-based technique we call Inverse Rational Control for inferring latent dynamics which could underlie rational thoughts. Then we offer a theoretical framework about neural coding that shows how to use these imputed rational thoughts to construct an interpretable description of neural dynamics.

We illustrate these contributions by analyzing a task performed by an artificial brain, showing how to test the hypothesis that a neural network has an implicit representation of task-relevant variables that can be used to interpret neural computation. As a case study, we choose a simple but ecologically critical task — foraging — whose solution requires an agent to account for the four crucial properties mentioned above: latent variables, partial observability, nonlinearities, and dynamics. Our general approaches should serve as valuable tools for interpreting behavior and brain activity for real agents performing naturalistic tasks.

## Results I: Modeling behavior as rational

In an uncertain and partially observable environment, animals learn to plan and act based on limited sensory information and subjective values. To better understand these natural behaviors and interpret their neural mechanisms, it would be beneficial to estimate the internal model and reward function that explains animals’ behavioral strategies. In this paper, we model animals as rational agents acting optimally to maximize their own subjective rewards, but under a family of possibly incorrect assumptions about the world. We then invert this model to infer the agent’s internal assumptions and rewards and estimate the dynamics of internal beliefs. We call this approach Inverse Rational Control (IRC), because we infer the reasons that explain an agent’s suboptimal behavior to control its environment.

This method creates a probabilistic model for an agent’s trajectory of observations and actions, and selects model parameters that maximize the likelihood of this trajectory. We make assumptions about the agent’s internal model, namely that it believes that it gets unreliable sensory observations about a world that evolves according to known stochastic dynamics. We assume that the agent’s actions are chosen to maximize its own subjectively expected long-term utility. This utility includes both benefits, such as food rewards, and costs, such as energy consumed by actions; it should also account for internal states describing motivation, like hunger or fatigue, that modulate the subjective utility. Finally, we assume that the agent follows a stationary policy based upon its mental model. This means that we cannot model learning with our method, although we can study adaptation and context dependence as long as our model represents these variables and their dynamics. We use the agent’s sequence of observations and actions to learn the parameters of this internal model for the world. Without a model, inferring both the rewards and latent dynamics is an underdetermined problem leading to many degenerate solutions. However, under reasonable model constraints, we demonstrate that the agent’s reward functions and assumed dynamics can be identified. Our learned parameters include the agent’s assumed stochastic dynamics of the world variables, the reliability of sensory observations about those world states, and subjective weights on action-dependent costs and state-dependent rewards.

### Partially Observable Markov Decision Process

To define the Inverse Rational Control problem, we first formalize the agent’s task as a Partially Observable Markov Decision Process (POMDP, Figure 1A) (10), a powerful framework for modeling agent behavior under uncertainty. A Markov chain is a temporal sequence of states *s* ∈ 𝒮 for which the transition probability *T* to the next state depends only on the current state, not on any earlier ones: *T* (*s*_*t*+1_ |*s*_0:*t*_) = *T* (*s*_*t*+1_| *s*_*t*_). A Markov Decision Process (MDP) is a Markov chain where at each time an agent can influence the world state transitions by deciding to take an action *a* ∈ 𝒜, according to *T* (*s*_*t*+1_ |*s*_*t*_, *a*_*t*_). At each time step the agent receives a reward or incurs a cost (negative reward) that depends on the world state and action, *R*(*s*_*t*_, *a*_*t*_). The agent’s goal is to choose actions that maximize its value *V*, measured by total expected future reward (negative cost) with a temporal discount factor *γ* ∈ (0, 1), so that 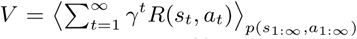, where the angle brackets denote an average with respect to the subscripted distribution. The actions are drawn from a state-dependent probability distribution called a policy, *π*(*a*|*s*_*t*_), which may be concentrated entirely on one action or may have some width. In a normal MDP, the agent can fully observe the current world state, but must plan for an unknown future. In a Partially Observed MDP (POMDP), the agent again does not know the future, but does not even know the current world state exactly. Instead the agent only gets unreliable observations *o* ∈ Ω about it, drawn from the distribution *o*_*t ∼*_*O*(*o*|*s*_*t*_). The agent’s goal is the same, to maximize the total expected temporally-discounted future reward. The POMDP is then a tuple of all of these mathematical objects: ℳ= (𝒮, *𝒜*, Ω, *R, T, O, γ*). Different POMDPs tuples reflect different tasks.

**Fig. 1.**
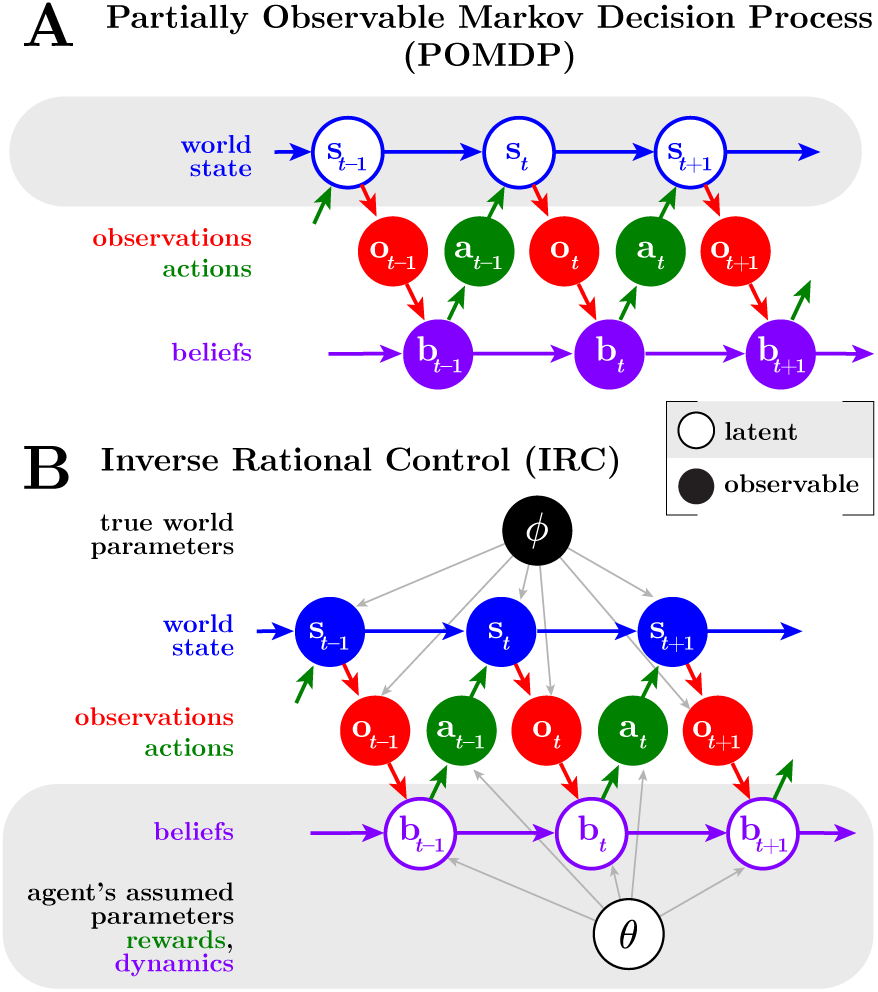
Graphical model of a Partially Observable Markov Decision Process (POMDP) (**A**) and the Inverse Rational Control (IRC) problem (**B**). Empty circles denote latent variables, and solid circles denote observable variables. For the POMDP, the agent knows its beliefs but must infer the world state. For IRC, the scientist knows the world state but must infer the beliefs. The real world dynamics depends on parameters *ϕ*, while the belief dynamics and actions of the agent depends on parameters *θ*, which include both its assumptions about the stochastic world dynamics and observations, and its own subjective rewards and costs.

Optimal solution of a POMDP requires the agent to compute a time-dependent posterior probability over the possible current world state *s*, given its history of observations and actions. Knowledge of all of that history can be summarized concisely in a single distribution, the posterior *B*(*s*). We consider this to reflect the *belief* of the agent about its current world state. It is useful to define a more compact *belief state b* as a set of sufficient statistics that completely summarize the posterior, so we can write *B*(*s*_*t*_|*b*_*t*_) = *B*(*s*_*t*_|*o*_1:*t*_, *a*_0:*t*−1_). This belief state can be expressed recursively using the Markov property as a function of its previous value (Supplemental Information Eq. 2).

We can express the entire partially observed MDP as a fully-observed MDP called a Belief MDP, where the relevant fully-observed state is not the world state *s* but instead the agent’s own belief state *b* (11). To do so, we must re-express the transitions and rewards as a function of these belief states (Supplemental Information, Eqs. 6,8). The optimal agent then determines a value function *Q*(*b, a*) over this belief space and allowed actions, based on its own subjective rewards and costs. This value can be computed recursively through the Bellman equation (12) (Supplemental Information, Eq. 9). The optimal policy deterministically selects whichever action maximizes the state-action value function. An alternative stochastic policy samples actions from a softmax function on value, 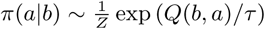 with a temperature parameter *τ* and normalization constant *Z*. The randomness introduces a new sub-optimality to the agent: instead of choosing the action with the maximal value, the agent has some chance of choosing a worse action. In the limit of a low temperature *τ* we recover the optimal policy, but a real agent may be better described by a stochastic policy with some controlled exploration.

### Inverse Rational Control

Despite the appeal of optimality, animals rarely appear optimal in experimentally defined tasks, and not just by exhibiting more randomness. Short of optimality, what principled guidance can we have about an animal’s actions that would help us understand its brain? One possibility is that an animal is ‘rational’ — that is, optimal for different circumstances than those being tested. In this section we present a behavioral analysis based on the possibility that agents are rational in this sense. The core idea is to parameterize possible strategies of an agent by those tasks under which each is optimal, and find which of those best explains the behavioral data.

We specify a *family* of POMDPs where each member has its own task dynamics, observation probabilities, and subjective rewards, together constituting a parameter vector *θ*. These different tasks yield a corresponding family of optimal agents, rather than a single optimized agent. We then define a log-likelihood over the tasks in this family, given the experimentally observed data and marginalized over the agent’s latent beliefs (Figure 1B):

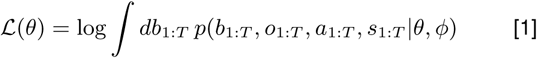

In other words, we find a likelihood over *which tasks* an agent solves optimally. In [1], *ϕ* are known parameters in the experimental setup that determine the world dynamics. Since they only affect observed quantities in the graphical model, they do not affect the model likelihood over *θ* (Supplementary Information).

This mathematical structure connects interpretable models directly to experimentally observable data, allowing us to formalize important scientific problems in behavioral neuroscience. For example, we can maximize the likelihood to find the best interpretable explanation of an animal’s behavior as rational within a model class, as we show below. We can also compare categorically different model classes that attribute to the agent different reward structures or assumptions about the task.

The log-likelihood [1] seems complicated, as it depends on the entire sequence of observations and actions and requires marginalization over latent beliefs. Nonetheless it can be calculated using the Markov property of the POMDP: the actions and observations constitute a Markov chain where the agent’s belief state is a hidden variable. We show that it is possible to exploit this structure to compute this likelihood efficiently (Supplemental Information).

### Challenges and solutions for rationalizing behavior

To solve the IRC problem, we need to parameterize the task, beliefs, and policies, and then we need to optimize the parameterized log-likelihood to find the best explanation of the data. This raises practical challenges that we need to address.

Our core idea for interpreting behavior is to parameterize everything in terms of tasks. All other elements of our models are ultimately referred back to these tasks. Consequently, the beliefs and transitions are distributions over latent task variables, the policy is expressed as a function of task parameters and preferences, and the log-likelihood is a function of the task parameters that we assume the agent assumes.

Thus, whatever representations we use for the belief space or policy, we need to be able to propagate our optimization over the task parameters through those representations. This is one requirement for practical solutions of IRC. A second requirement is that we can actually compute the optimal policies.

Efficient representation of general beliefs and transitions is hard since the space of probabilities is much larger than the state space it measures. The belief state is a probability distribution and thus takes on continuous values even for discrete world states. For continuous variables the space of probabilities is potentially infinite-dimensional. This poses a substantial challenge both for machine learning and for the brain, and finding neurally plausible representations of uncertainty is an active topic of research (13–18). We consider two simple methods to solve IRC using lossy compression of the beliefs: discretization, or distributional approximation. We then provide a concrete example application in the discrete case.

#### Discrete beliefs and actions

If we have a discrete state space then we can use conventional solution strategies for Markov Decision Processes. For a small enough world space, we can exhaustively discretize the complete belief space, and then solve the Belief MDP problem with standard MDP algorithms (12, 19). In particular, the state-action value function *Q*(*b, a*) under a softmax policy *π*(*a*|*b*) can be expressed recursively by a Bellman equation, which we solve using value iteration (11, 12). The resultant value function then determines the softmax policy *π*, and thereby determines the policy-dependent term in the log-likelihood [1].

Finally, to solve the IRC problem we can directly optimize this log-likelihood, for example by greedy line search (Supplementary Information). An alternative in higher-dimensional problems is to use Expectation-Maximization to find a local optimum, with a gradient ascent M-step (Supplementary Information, (20, 21)). To compute the gradient of the log-likelihood, we again use recursion to calculate the value gradient *∂Q/∂θ* exactly, and use the chain rule to derive the policy gradient and then the 𝒬 auxiliary function gradient (Supplementary Information).

#### Continuous beliefs and actions

The computational expense of the discrete solution grows rapidly with problem size, and become intractable for continuous state spaces and continuous controls. A practical choice is to continually update a finite set of summary statistics as for an extended Kalman filter, although it may be tractable to learn and use a more general set of statistics (18). Rational control with continuous actions also requires us to implement a flexible family of continuous policies *π* that map from beliefs to actions. We use deep neural networks to implement these policies (22). Deep learning methods are commonly used in reinforcement learning to provide flexibility, but they lack interpretability: information about the policy is distributed across the weights and biases of the network. Crucially, to maintain interpretability, we parameterize this family by the task. Specifically, we provide the model parameters as *additional inputs* to a policy network, and learn the optimal policies simultaneously over a prior distribution on task parameters *p*(*θ*) (22). This allows the network to generalize its optimal strategies across POMDPs in the task family. It also allows us to easily maximize the likelihood (Eq 1) by gradient ascent using auto-differentiation (22).

### Application to foraging

We applied our analyses to understand the workings of a neural network performing a foraging task. The task requires an agent to combine unreliable sensory data with an internal memory to infer when and where rewards are available, and how to best acquire them. We train an artificial recurrent neural network to solve this task in a suboptimal but rational way and use Inverse Rational Control to infer its assumptions, subjective preferences, and beliefs.

#### Task description

Two locations (‘feeding boxes’) have hidden food rewards that appear and disappear according to independent telegraph processes with specified transition probabilities (Figure 2, (23)). The boxes provide unreliable color cues about the current reward availability, ranging from blue (probably unavailable) to red (probably available). We assume there are three possible locations for the agent: the locations of boxes 1 and 2, and a middle location 0. We include a small ‘grooming’ reward for staying at the middle location, to allow the agent to stop and rest. A few discrete actions are available to the agent: it can push a button to open a box to either get reward or observe its absence, it can move toward a new location, or it can do nothing. Traveling and pushing a button to open the box each have an associated cost. This disincentivizes the agent from repeating fruitless actions. When a button-press action is taken to open a box, any available reward there is acquired. Afterwards, the animal knows there is no more food available now in the box (since it was either unavailable or consumed) and the belief about food availability in that box is reset to zero. The specific values of these parameters used in our experiments are described in Supplemental Information.

**Fig. 2.**
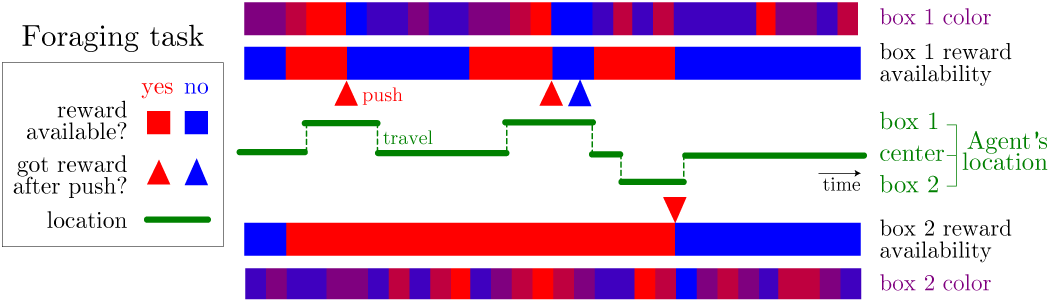
Illustration of foraging task with latent dynamics and partially observable sensory data. The reward availability in each of two boxes evolves according to a telegraph process, switching between available (red) and unavailable (blue), and colors give the animal an ambiguous sensory cue about the reward availability. The agent may travel between the locations of the two boxes. When a button is pressed to open a box, the agent receives any available reward.

#### Neural network agent

To test the IRC algorithm and our subsequent neural coding analyses, we wanted a synthetic brain for which we could assess the ground truth. We therefore used imitation learning to train a neural network to simultaneously reproduce the policies of multiple rational agents. These rational agent “teachers” each optimally solve one POMDP problem in our task family (Methods). Figure S1A shows the architecture of our recurrent network. The network receives sensory inputs about the location, color cues, and rewards, plus additional inputs specifying the task parameters. After training to match the rational agents, outputs of the neural network closely match the POMDP agent’s policies (Figure S1B), but these task-relevant quantities are encoded implicitly in a large population of neurons.

Finally, to impose suboptimality upon our neural network agent, we misled it by providing inputs that specified the wrong set of task parameters, while testing the agent on a task with different parameters. These inputs led to a time series of observations *o*_*t*_, actions *a*_*t*_, and neural activity ***r***_*t*_. Together these constitute the experimental measurements for our suboptimal agent.

#### Inverse Rational Control for foraging

In our target applications, we don’t know the agent’s assumed world parameters, nor their subjective costs, nor the amount of randomness (softmax temperature). Our goal is to estimate a simulated agent’s internal model and belief dynamics from its chosen actions in response to its sensory observations. We infer all of these using IRC.

The actions and sensory evidence (color cues, locations and rewards) obtained by the agent all constitute observations for the experimenter’s learning of the agent’s internal model. Based on these observations over 5000 time points, including 1595 movements and 566 button presses, we use IRC to infer the parameters of the internal model that can best explain the behavioral data (Figure 3A). Figure 3B shows that IRC correctly imputes a rational model to the neural network, whose parameters closely match those of its teacher.

**Fig. 3.**
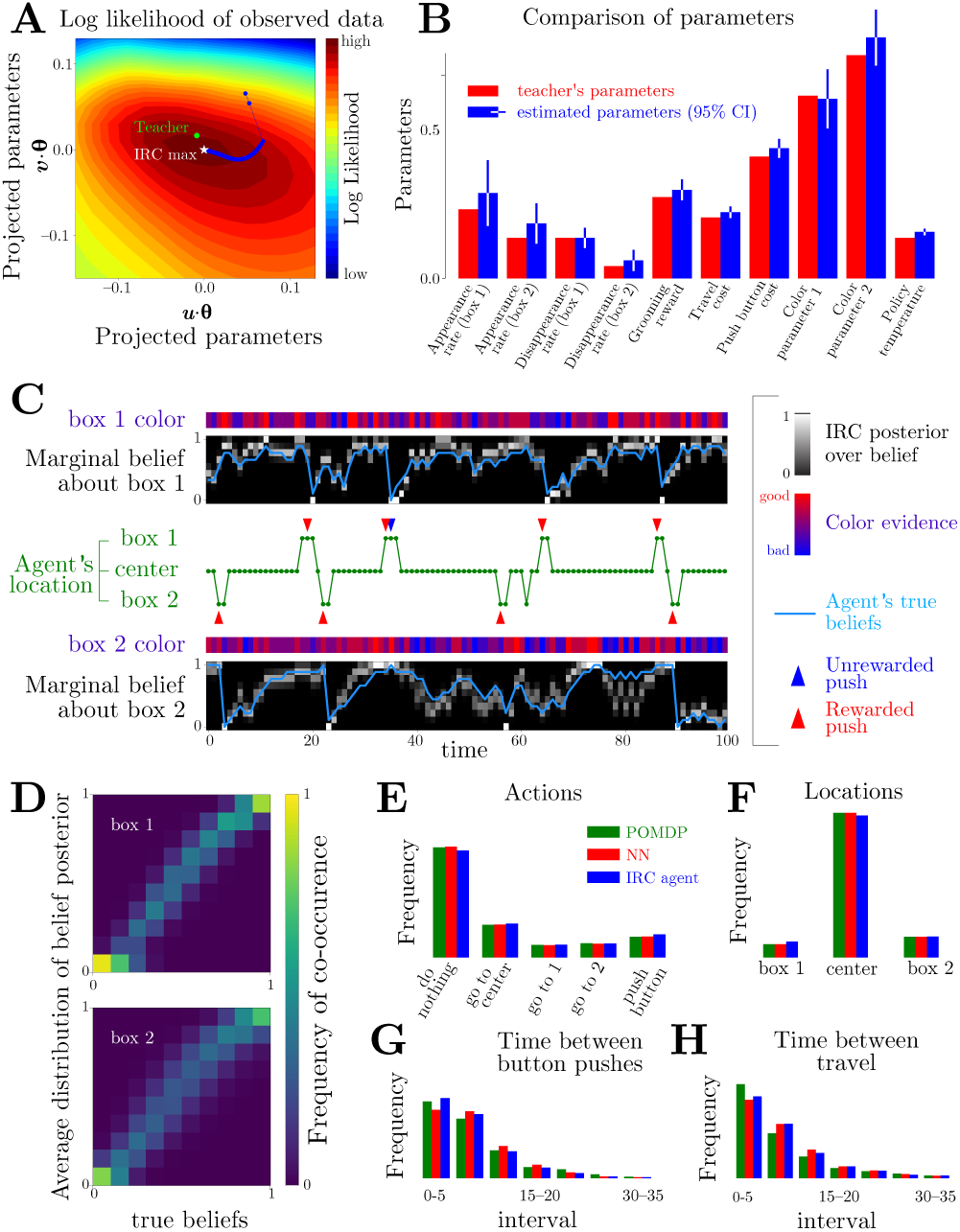
Successful recovery of agent model by Inverse Rational Control. The agent was a neural network trained to imitate a suboptimal but rational teacher. (**A**) The estimated parameters converge to the optimal point of the observed data log-likelihood (white star). Since the parameter space is high dimensional, we project it onto the first two principal components ***u, v*** of the learning trajectory for *θ* (blue). (**B**) Comparison of the true parameters of the agent and the estimated parameters. Error bars show 95% confidence intervals based on the Hessian of log-likelihood (Supplemental Information Figure S2). (**C**) Estimated and true marginal belief dynamics over latent reward availability. These estimates are informed by the noisy color data at each box and the times and locations of the agent’s actions. The posteriors over beliefs are consistent with the dynamics of the true beliefs (blue line). (**D**) The mean posterior beliefs, 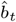, 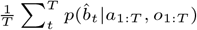 closely match the true beliefs of the teacher. Inferred distributions of (**E**) actions, (**F**) residence times, (**G**) intervals between consecutive button-presses, and (**H**) intervals between movements.

Data limitations imply some discrepancy between the true parameters and the estimated parameters which can be reduced with more data. With the estimated parameters, we are able to infer a posterior over the dynamic beliefs (Figure 3C). (Note that this is an experimenter’s posterior over the agent’s subjective posterior!) Although we do not know what the neural network believes, the inferred posterior is consistent with the imitated teacher’s subjective probabilities of food availability in each box. The inferred distributions over beliefs reveals strong correlations between the belief states of the teacher and the belief states imputed to the neural network (Figure 3D).

Figure 3E–H shows that the artificial brain and inferred agent choose actions with similar frequencies, occupy the three locations for the same fraction of time, and wait similar amounts of time between pushing buttons or travelling. This demonstrates that the IRC-derived agent’s internal model generates behaviors that are consistent with behaviors of the agent from which it learned.

## Results II: Neural coding

We don’t presume that any real brain explicitly calculates a solution to the Bellman equation, but rather learns a policy by combining experience and mental modeling. With enough training, the result is an agent that behaves ‘as if’ it were solving the POMDP (Figure 4A). In this section we present a framework for understanding the brain computations that could implement such behaviors.

**Fig. 4.**
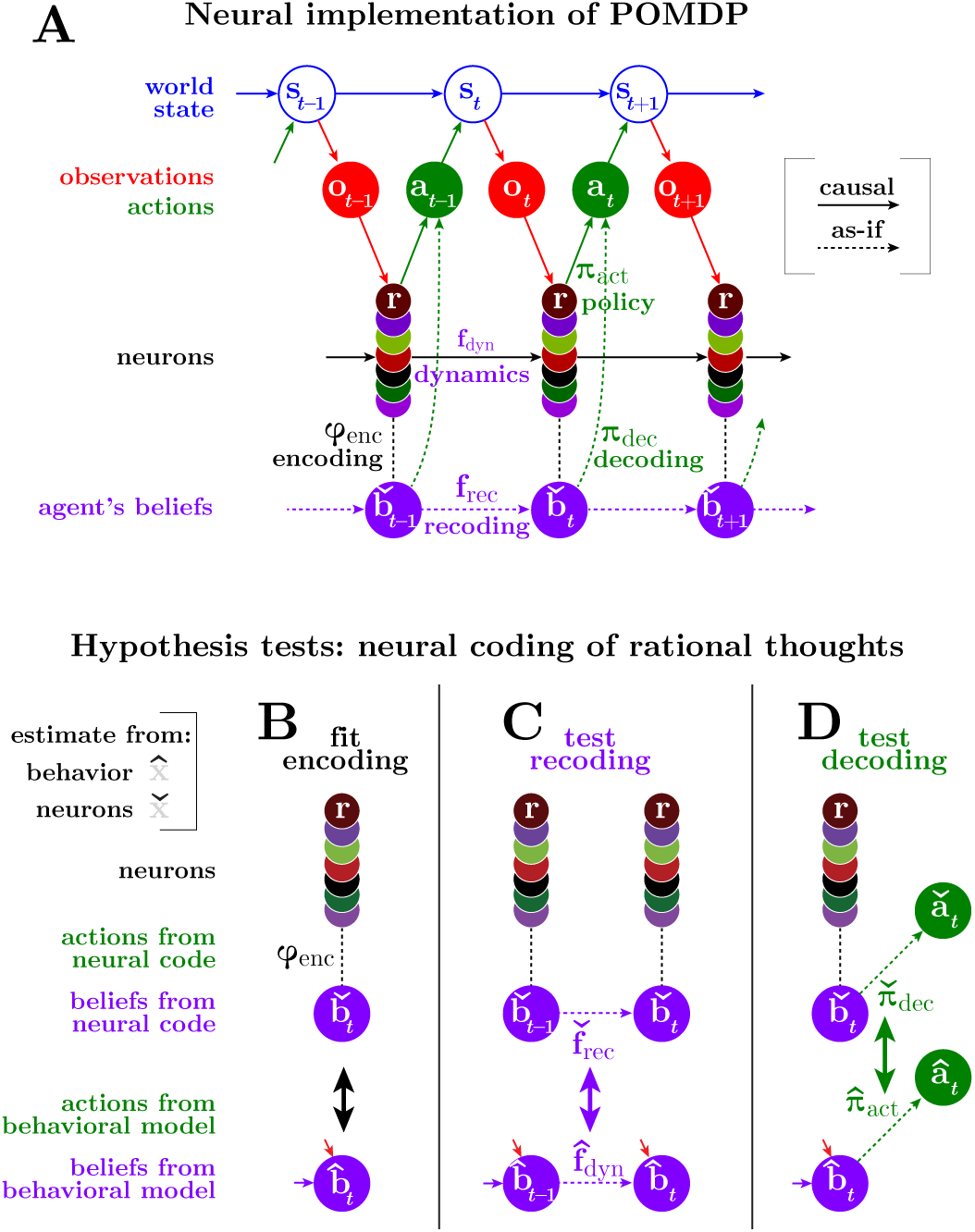
Schematic for analyzing a dynamic neural code. (**A**) Graphical model of a POMDP problem with a solution implemented by neurons implicitly encoding beliefs. We find how behaviorally relevant variables (here, beliefs) are *encoded* in measured neural activity through the function 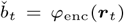 (**C**) We then test our hypothesis that the brain *recodes* its beliefs rationally by testing whether the dynamics of the behaviorally estimated belief 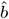 match the dynamics of the neurally estimated beliefs 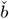, as expressed through the update dynamics 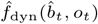 and recoding function 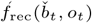. (**D**) Similarly, we test whether the brain *decodes* its beliefs rationally by comparing the behaviorally and neurally derived policies 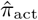 and 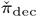. Quantities estimated from behavior or from neurons are denoted by up-pointing or down-pointing hats, ^ and ̌(Table S1).

Many researchers in machine learning express skepticism that we can find much that is human-interpretable about either artificial or biological neural networks (24, 25). One interesting counter-argument is that near any solution found by machine learning optimization, there may be other solutions that perform similarly while retaining interpretability (26). Additionally, solutions that match the causal structure of the environment are naturally more interpretable and tend to generalize better (27, 28), and thus may be favored by biological learning. More humbly, even if we cannot find an interpretable network that exactly instantiates the brain’s computations, we may still glean satisfying and useful insights from partial explanations at a higher level of abstraction (29–31).

To move toward more interpretable computations, our analysis does not focus on neural responses, but rather on the task-relevant information encoded in those responses. Targeted dimensionality reduction abstracts away the fine details of the neural signals in favor of an algorithmic- or representational-level description. This can decrease the number of parameters needed to characterize dynamics, reducing overfitting. More importantly, it can avoid the massive degeneracies inherent in neuron-level mechanisms: different neural networks could have entirely different neural dynamics but could share the task-relevant computations. This indicates how a deeper, more invariant understanding of neural computations is possible at the algorithmic level than at the mechanistic level (32).

Analysis of the linked processes of encoding, recoding, and decoding can help interpret task-relevant computations. These processes correspond to representation, dynamics, and action. The brain’s ‘encoding’ specifies the task-relevant and -irrelevant coordinates of neural activity (Figure 4B). ‘Recoding’ describes how that encoding is transformed over time and space by neural processing (Figure 4C). ‘Decoding’ describes how those estimates predict future actions (Figure 4D).*

The neural coding framework makes one crucial assumption: the neural data must be sufficient to capture the task-relevant neural processes. The important aspects are different for encoding, recoding, and decoding. To describe the *encoding*, we need to measure the right neurons at the right resolution to be sensitive to the task-relevant properties, which may include nonlinear statistics (33, 34) and will certainly exhibit some variability (35). To describe the *recoding* accurately, all measured changes in neural state must depend only on the current state. In other words, the measured neural dynamics should be Markovian. Markovian dynamics are an essential property of any causal system. To describe *decoding* accurately, we must measure the neural signals that eventually drive the behavior. If the chosen state space lacks any of this relevant information due to missing neurons, slow measurements, lossy post-processing, *etc.*, then we will see unexplainable variability in the encoded variables, recoding dynamics, and decoded actions. As long as we do measure the right signals, our neural coding framework applies equally well to spiking, multi-unit activity, calcium concentration, neurotransmitter concentration, LFPs, conventional frequency bands, or any other signals hypothesized to contain task-relevant information. For example, if distinct neural frequency bands encode distinct information or interactions, then slow firing rates alone will not be sufficient to capture dynamics. Nonetheless, in such cases we may be able to construct a sufficient state space by augmenting the neural states, for example by explicitly including multiple frequency bands or the recent firing rate history.

Once we fit a neural encoding, we subsequently concentrate only on the task-relevant coordinates specified by that encoding. By construction, this level of explanation need not capture every facet of neural responses nor the physical mechanism by which they evolve. Obviously it cannot explain responses to untested task variables. Nonetheless, it would be great progress if we can account for stimulus- and action-dependent neural dynamics within a task-relevant coordinates (36) that explains how pieces of information interact and predict behavior. Although this ‘as-if’ description cannot legitimately claim to be causal, it can be promoted to a causal description since it does provide useful predictions for causal tests about what neural features should influence computation and action (37, 38).

Just as a complete description of neural mechanisms requires those dynamics to be Markovian, a complete lower-dimensional description of task-relevant computations also requires that the dynamics are Markovian. In other words, we seek task-relevant coordinates whose updates depend only on those coordinates. Otherwise we will again find unexplained variability in the task-relevant dynamics (Figure S3) or actions.

Figure 5 provides a conceptual illustration of the geometry of task-relevant and -irrelevant coordinates in neural activity space, and the types of errors that can occur when measuring task-relevant neural computation. Neural activity occupies a manifold of much lower dimension than the ambient space of all possible neural responses (39). Within that manifold there is further structure, with task-relevant variables tracing out submanifolds related to each other by task-irrelevant neural variations.

**Fig. 5.**
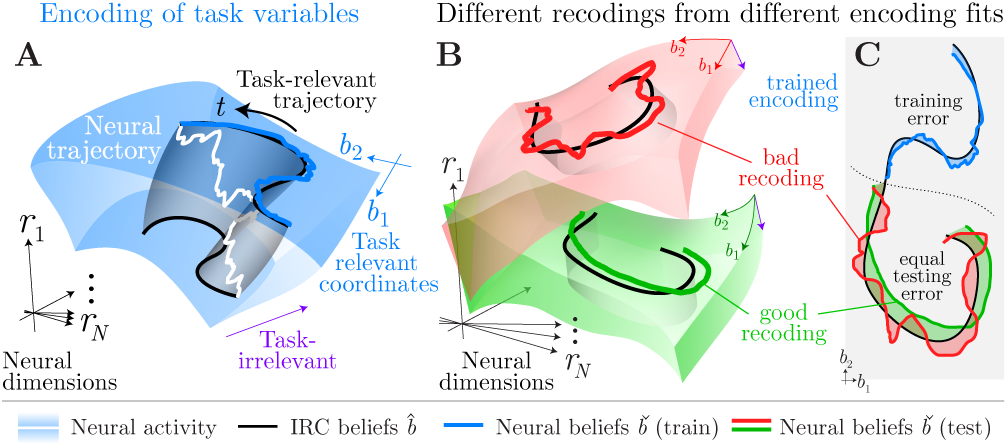
Conceptual illustration of encoding and recoding. (**A**) Neural responses ***r*** inhabit a manifold (blue volume, here three-dimensional) embedded in the high-dimensional space of all possible neural responses. A neural encoding model divides this manifold into task-relevant and -irrelevant coordinates (blue and purple axes). We must estimate these coordinates from training data, given some inferred task-relevant targets ***b***. According to this encoding, many activity patterns ***r*** can correspond to the same vector of task variables ***b***. Any particular neural trajectory (white curve) is just one of many that would trace out the same task-relevant projection ***b***(*t*) (black curves). The set of all neural activities consistent with one task-relevant trajectory therefore sweeps out a manifold (grey ribbon). (**B**) After fitting an estimator of the task variables using *training* data, we can measure how well the encoding describes the task variables in a new *testing* data set. Different encodings (red and green volumes) divide the same neural manifold differently into relevant and irrelevant coordinates, and the task variables **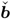** estimated from these neural encodings (red and green curves) will deviate in different ways from the variables **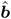** inferred from behavior (black). (**C**) The testing error of these neurally derived task variables (red, green) will be larger than the training error (blue). Task-relevant variables **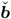** derived from different encoding models may have the same total errors, but may nonetheless have different recoding dynamics. Here the smoother green dynamics are closer to the behaviorally inferred dynamics than the rougher red dynamics, which implies that these task-relevant dimensions better capture the computations implied by Inverse Rational Control. Supplemental Figure S3 provides more detail of good and bad recodings.

In principle, our framework can apply to many different tasks and computations. For concreteness, here we will present our analysis using the computations and variables inferred by Inverse Rational Control. The inferred internal model allows us to impute the agent’s time-dependent beliefs *b* about the partially observed world state ***s***. Such a belief vector might include full posterior over the world state, *B*(*s*_*t*_ |*o*_1:*t*_, *a*_1:*t* −1_) as we used for the discrete IRC above, or a point estimate *ŝ* of the world state and a measure of uncertainty about it, say a covariance Σ_*s*_, as in the Gaussian approximation we have used for continuous IRC (22). To us, as scientists, the agent’s beliefs are latent variables, so our algorithm can at best create a posterior *p*(*b*) over those beliefs, or a point estimate 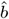 indicating the most probable belief. Here we will base our analyses on a point estimate over beliefs. Below we describe our general analysis approach and apply it to understand the neural computations implemented during foraging by the simulated brain.

### Encoding

Given beliefs 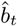 imputed by IRC, we can estimate how they are encoded in the neural responses ***r***. An encoding defines a response distribution 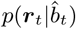, which determines both task-relevant and -irrelevant coordinates (Figure 5A). To find what is encoded by this probabilistic mapping, we use a (potentially nonlinear, potentially even spatiotemporal) readout function *ϕ*_enc_(***r***_*t*_) fit to minimize the discrepancy between the behavioral target belief 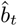 and the neural estimate 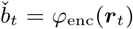 (Figure 4B). After training *ϕ*_enc_ to match the behavioral targets 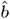 and ignore task-irrelevant aspects of the neural responses, we can then cross-validate it on new estimates 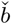 from fresh neural data. Since data is finite and noisy, the models invariably have some errors caused by deviations between the estimated task-relevant coordinates and the true ones. These errors are smaller for the training data, and larger for fresh testing data. Different fits of encoding models partition the neural manifold differently, and will thus generally have different testing errors (Figure 5B).^†^

### Recoding

Recoding describes the changes in a neural encoding. While neural dynamics may affect every dimension of neural activity, we focus only on the low-dimensional, interpretable dynamics within the neural manifold. By construction, those dynamics reflect the changes in the agent’s beliefs.

The rational control model predicts that beliefs are updated by sensory observations and past beliefs, with interactions that are determined by the internal model according to a function *b*_*t*+1_ = *f*_dyn_(*b*_*t*_, *o*_*t*_) + *η*_*t*_ where *f*_dyn_ and *η*_*t*_ reflect the task-relevant and -irrelevant parts of the dynamics (the latter may include stochastic components as well as deterministic components that depend on task-irrelevant dimensions, Figures 5 and S3). If our neural analysis correctly identifies dynamics responsible for behavior, then the beliefs 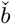 estimated from the neural encoding should be recoded over time following those same update rules. We estimate this neural recoding function 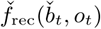 directly from the sequence of neurally estimated beliefs 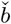 by minimizing differences between the actual and predicted future neural beliefs. We then compare 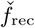 to the update dynamics posited by the behavioral model 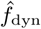 (Figure 4C). (Note that we should compare these only over the distribution of experienced beliefs, *i.e.* those beliefs for which the recoding function matters in practice.) Agreement between the behavioral belief dynamics and the neurally derived belief dynamics implies that we have successfully understood the ‘recoding’ process. Even for good encoding models this is not guaranteed, since activity outside the encoding manifold could influence the neural dynamics: Two different fitted encoding models could provide equal reconstruction errors, and yet because of limited data or model mismatch only one has neural dynamics that match the behaviorally derived dynamics (Figure 5B,C).

### Decoding

These encodings and recodings do not matter if the brain never decodes that information into behavior. We can evaluate how the brain uses its information by fitting a policy 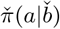 to predict observed actions directly from the neurally encoded beliefs. We then test the hypothesis that the brain decodes neurally encoded rational thoughts by comparing that neurally-derived policy 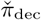 against the behavioral policy, 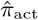 (Figure 4D).

### Application to simulated foraging agent

Figure 6 presents the results of applying this neural coding framework to look inside the brain of our simulated agent while it forages.

**Fig. 6.**
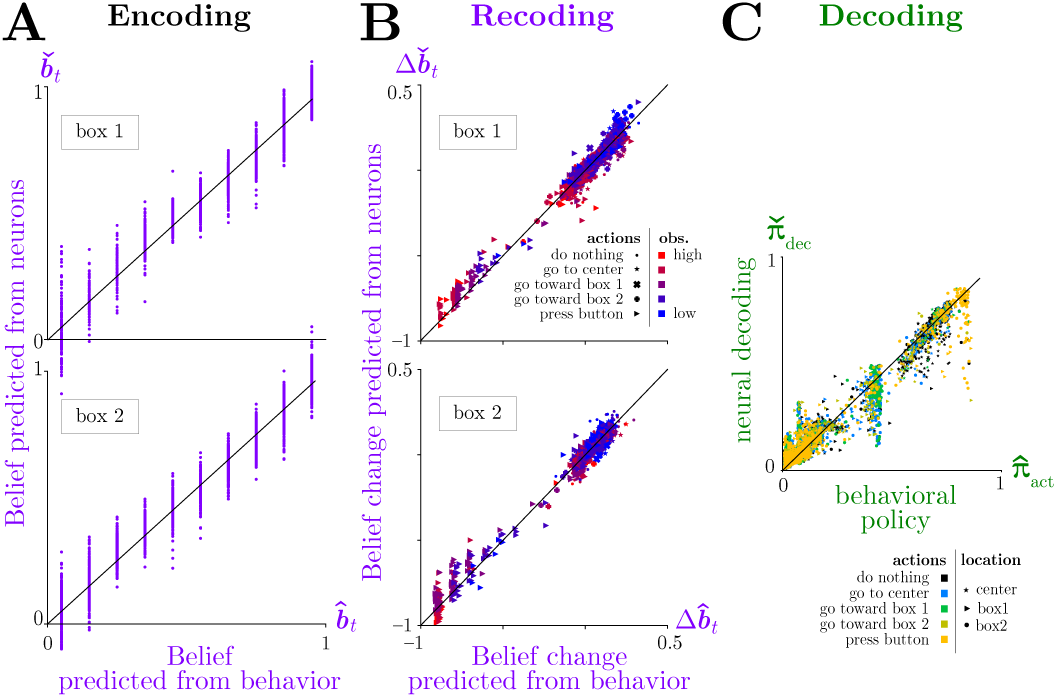
Analysis of neural coding of rational thoughts. (**A**) Encoding: Neurally-derived beliefs 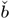 match behaviorally-derived beliefs 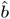 based on IRC. Cross-validated neural beliefs are estimated from testing neural responses ***r*** using a linear estimator, 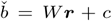, with weight matrix fit from separate training data. (**B**) Recoding: Belief updates 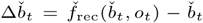 from the neural recoding function 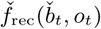 match the corresponding belief updates from the task dynamics 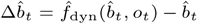. Neural updates are estimated using nonlinear regression with radial basis functions (Methods). (**C**) Decoding: The policy 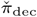 predicted by decoding neural beliefs approximately matches the policy 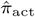 estimated from behavior by IRC. Neural policy is estimated from actions *a* and neural beliefs 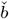 using nonlinear multinomial regression (Methods).

To evaluate the encoding for our synthetic brain, we assume that beliefs *b*_*t*_ are linearly encoded instantaneously in neural activity ***r***_*t*_. After performing linear regression of behaviorally derived beliefs 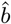 against neural activity ***r***, we can estimate other beliefs 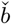 from previously unseen neural data. Figure 6A shows that these beliefs estimated from neural data are accurate.

Figure 6B shows that the recoding dynamics obtained from the neural belief dynamics also match the dynamics described by the rational model. We characterize these neural dynamics using kernel ridge regression between 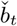 and 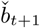 (Methods). The resultant temporal changes in the neurally-derived beliefs 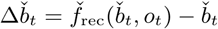 agree with the corresponding changes in the behavioral model beliefs, 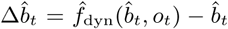. Although some of these changes are driven directly by the sensory observations (colors), that only explains part of the belief updates: even conditioned on a given sensory input at one time, the updates agree between the neurons and the behavioral model. This provides evidence that we understand the internal model that governs recoding at the algorithmic level.

To account for the discrete actions space, our example analysis of neural decoding uses nonlinear multinomial regression to fit the probabilities 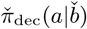 of allowed actions as a function of neurally derived beliefs (Methods). Comparing the resultant function to the rational policy 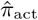 shows that these two decoding functions match well (Figure 6C). This provides evidence that we understand the decoding process by which task-relevant neural activity generates behavior.

## Discussion

This presents an explainable AI paradigm to infer an internal model, latent beliefs, and subjective preferences of a rational agent that solves a complex dynamic task described as a Partially Observable Markov Decision Process. We fit the model by maximizing the likelihood of the agent’s sensory observations and actions over a family of tasks. We then described a neural coding framework for testing whether the imputed latent beliefs encoded in a low-dimensional manifold of neural responses are recoded and decoded in a manner consistent with this behavioral model. We demonstrated these two contributions by analyzing the neural coding of an implicit computational model by an artificial neural network trained to solve a simple foraging task requiring memory, evidence integration, and planning. For this simulated data, we successfully recovered the agent’s internal model and subjective preferences, and found neural computations consistent with that model.

### Related work

Our approach generalizes previous work in artificial intelligence on the inverse problem of learning agents by observing behavior. Methodologically, other studies of inverse problems address parts of Inverse Rational Control, but with a non-scientific goal — getting artificial agents to solve tasks by learning from demonstrations of expert behavior. Inverse Reinforcement Learning (IRL) tackles the problem of learning how an agent judges rewards and costs based on observed actions (40), but assumes a known dynamics model (21, 41). Conversely, Inverse Optimal Control (IOC) learns the agent’s internal model for the world dynamics (42) and observations (43), but assumes the reward functions. In (44, 45) both reward function and dynamics were learned, but only the fully-observed MDP case is explored. We solve the natural but more difficult partially-observed setting, and ensure these solutions provide a scientific basis for interpreting animal behavior.

As a cognitive theory, by positing a rational but possibly mistaken agent, our approach resembles Bayesian Theory of Mind (BToM) (46–51). Previous work in BToM has considered tasks with uncertainty about static latent variables that were unknown until fully observed (51), or tasks with partially observed variables but simpler trial-based structure (46, 47). Here we allow for a more natural world, with dynamic latent variables and partial observability, and we infer models where agents make long-term plans and choose sequences of actions. Where prior work in BToM learned subjective rewards (51) *or* internal models (49), our Inverse Rational Control infers both internal models *and* subjective preferences in a partially observable world.

In addition, BToM studies have focused their attention on models of behavior, whereas our purpose is to connect dynamic model computations to brain dynamics. Some work has posited a POMDP model for behavior and hypothesized how specific brain regions might implement the relevant computations (52). Here we demonstrate an analysis framework to test such connections, by examining neural representations of latent variables and showing how computational functions could be embodied by low-dimensional neural dynamics.

While low-dimensional neural dynamics is an important topic for emerging studies of large-scale neural activity (2, 6, 7, 39), few have been able to relate these dynamic activity patterns to interpretable latent model variables. Far more commonly, these low-dimensional manifolds are attributed to an intrinsically generated manifold (36, 53), or are related to measurable quantities like sensory inputs or behavioral outputs (2, 54, 55). Population activity in the visual system is known to relate to latent representations extracted by trained deep networks (3, 4), and while this shows that many task-relevant features extracted by machine learning solutions are also task-relevant for the visual system, these feature sets yet account for neither temporal dynamics nor uncertainty, nor are they readily interpretable (56). Our proposed model-based analysis of population activity is currently our best bet for finding interpretable computational principles.

### Limitations and Generalizations

We demonstrated our approach to understanding cognition and neural computation by applying it to a task involving multiple important features, namely partially observable latent variables with structured dynamics requiring nonlinear computation. However, this foraging task is still fairly simple. Our conceptual framework is much more general and should be able to scale to more complex tasks. As we showed, it can model common errors of cognitive systems, such as inferring false beliefs derived from incorrect or incomplete knowledge of task parameters. But it can also be used to infer incorrect *structure* within a given model class. For example, it is natural for animals to assume that some aspects of the world, such as reward rates at different locations, are not fixed, even if an experiment actually uses fixed rates (57). Similarly, an agent may have a superstition that different reward sources are correlated even when they are independent in reality. Given a model class that includes such counterfactual relationships between task variables, our method can test whether an agent holds these incorrect assumptions. Our framework can also be generalized to cases of bounded rationality (58) by incorporating additional internal representational or computational constraints, such as metabolic costs (59) or architectural constraints (60). However, our approach does use model-based reinforcement learning and thus does require a model. Like any model-based algorithm, it can only explain behaviors we can represent by the policies that the model can generate. Moreover, even if the model can represent these policies in principle, it must be able to learn that family of policies. This can pose practical challenges that modern reinforcement learning methods are making rapid progress in overcoming.

Our core assumption for the behavioral model is that animals assume the world is Markovian, which leads them to use stationary policies. What if they don’t, due to a changing task or motivation? By including additional latent states, such as slow context variables or an internal motivation state, we may recover a stationary policy, and then our approach is again applicable. That said, this will be a poor model while the animal is learning something for the first time, and a higher-level rational learning model will be required.

There is a trade-off between model interpretability and flexibility. Large-scale tasks are now being solved with expressive neural networks (61, 62) that provide rich state representations, but may not permit interpretation. This may be an unavoidable limitation in a world of complex structure (24, 25). Or it may be that these uninterpretable representations are insufficiently constrained, and that richer tasks, multi-task training, and priors favoring sparse causal interactions may bias networks toward more human-interpretable representations (26, 28, 63, 64) that relate more closely to actionable latent variables (65).

When there is insufficient data to distinguish possible rational models, we may recover a sloppy model (66, 67) for which multiple combinations of parameters have the same likelihood (Eq. 1). We evaluated this in our foraging application by computing the curvature of the observed data log-likelihood (Figure S2), and found that our models were sufficiently constrained that all parameters were identifiable, although some combinations produced more optimistic beliefs compensated by higher action costs to generate similar action sequences.

### Conclusion

The success of our methods on simulated agents suggests it could be fruitfully applied to experimental data from real animals performing such foraging tasks (23, 68), as well as to richer tasks requiring even more sophisticated computations. Using explainable AI to construct belief states, their dynamics, and their utility for solving interesting tasks will provide useful targets for interpreting dynamic neural activity patterns, which could help identity the neural substrates of thoughts.

## Materials and Methods

### Inverse Rational Control

Full mathematical details for IRC, foraging task, and neural network training are available in Supplementary Information. Parameters were selected to expose interesting behaviors, such as balancing the relevance of predictable dynamics with sensory cues, and ensuring that both foraging sites were visited. Code for the discrete case is available at https://github.com/XaqLab/IRC_TwoSiteForaging.

### Neural coding analysis

#### Encoding

We find an encoding matrix 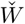 by regressing *b* against ***r***. This produces neural estimates of task-relevant variables 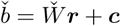 for new data. *Recoding*: We find dynamics by regressing 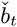 against 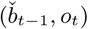 with kernel ridge regression. The kernel functions are radial basis functions with centers on all possible target beliefs and a width at half-max equal to the spacing between discrete beliefs. This yields the ‘recoding’ function 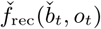 representing the nonlinear dynamics of the neural beliefs. We compare the belief updates 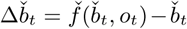 from the recoding function 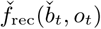 and the corresponding belief updates from the task dynamics 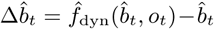. *Decoding*: We compute the brain’s ‘decoding’ function, *i.e.* an approximate policy 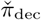, using nonlinear multinomial regression of 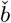 against *a* with the same radial basis functions as used in recoding. We use a feature space of radial basis functions with centers on a 9 × 9 grid over beliefs, with width equal to the center spacing, and an outer product space over locations.

## Acknowledgments

The authors thank Dora Angelaki, Baptiste Caziot, Valentin Dragoi, Krešimir Josić, Zhe Li, Rajkumar Raju, and Neda Shahidi for useful discussions. ZW, PS, and XP were supported in part by BRAIN Initiative grant NIH 5U01NS094368. ZW and XP were supported in part by an award from the McNair Foundation. SD and XP were supported in part by the Simons Collaboration on the Global Brain award 324143 and NSF 1450923 BRAIN 43092. XP and MK were supported in part by NSF CAREER Award IOS-1552868.

---

**Table S1.**
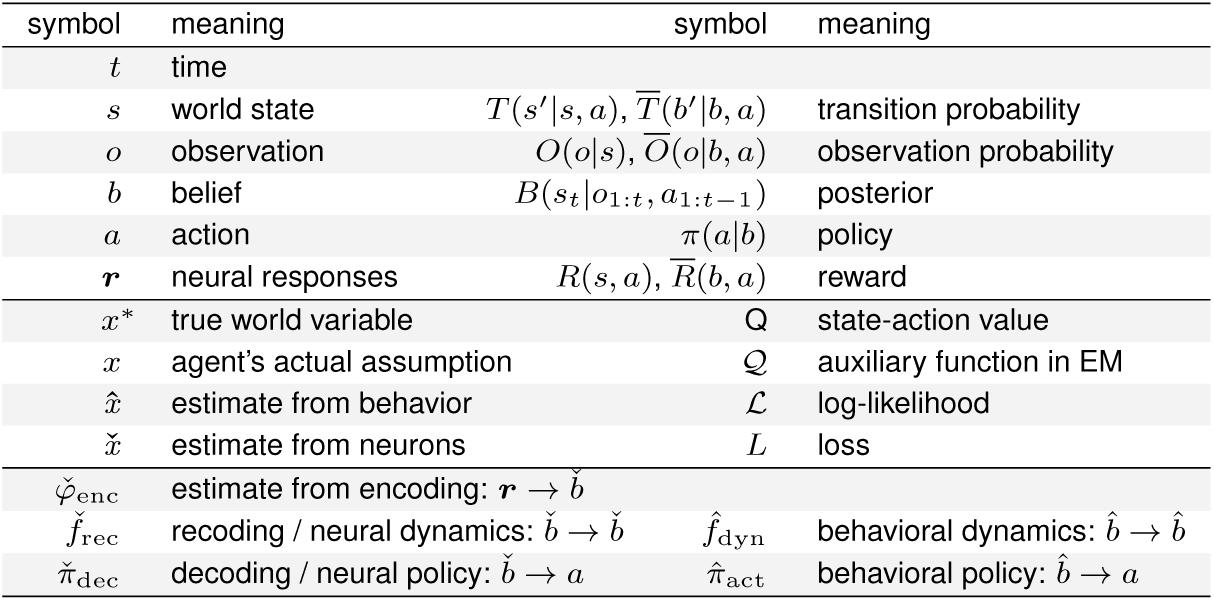
Glossary of notation.

## Supporting Information Appendix (SI)

### Belief MDP

In a belief MDP, an agent chooses actions based on the belief state *b*_*t*_, so the agent must compute the belief state at each time given its observations and actions up to that time. This can be computed online using the Markov property, according to

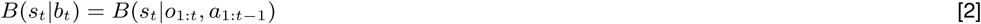

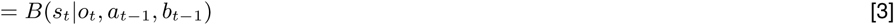

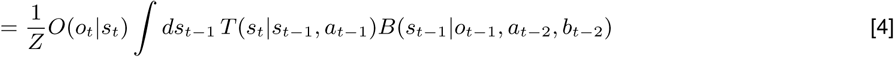

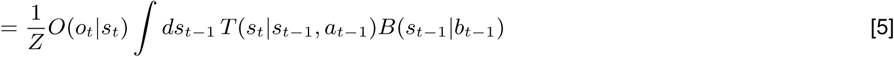

One subtle point to note is that for a deterministic policy, for which *π*(*a*|*b*) = *δ*(*a* − *a*\*(*b*)) with optimal action *a*\*(*B*), the next belief state is only a function of the last belief state and the current observation. Although the next world state does of course depend on the selected action, this action is perfectly predictable from the last belief state and so *P* (*b*_*t*_ + 1 |*b*_*t*_, *a*_*t*_, *o*_*t*_) = *P* (*b*_*t*_ + 1|*b*_*t*_, *o*_*t*_). In contrast, for a stochastic policy like the one we allow in our simulations, the belief state does not fully determine the action, and thus to know future belief states the agent would have to also observe the action that was actually sampled from the policy. This could be notated in multiple equivalent ways. For example, the graphical models in Figure 1 could include arrows from each action to the next belief state. Alternatively, the selected action could be considered part of the world state that is perfectly observed, and so it would be transmitted automatically to the next belief state without the addition of any explicit arrows. For ease of exposition and clarity of the trellis diagram we have chosen this latter approach. The mathematics, however, make these particular dependencies explicit.

To find the optimal policy, an agent evaluates the value of each action and state. If the agent were given future observations and actions, then its future beliefs would be known. But when observations are unknown, the agent has only a distribution over beliefs, arising from the distribution of future observations it may encounter from the distribution of future world states. The transition probability between belief states is then

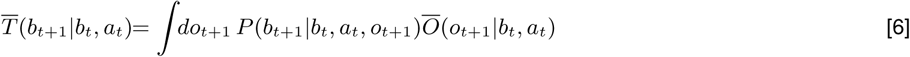

where

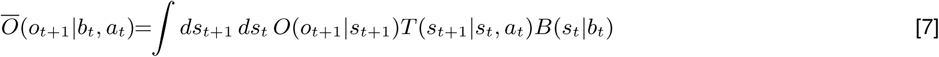

is the distribution of future observations given the present belief and action. The parameters of this belief transition probability 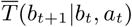 therefore include parameters from both the world state transitions *T* (*s*_*t*+1_ |*s*_*t*_, *a*_*t*_) and observation functions *O*(*o*_*t*_ |*s*_*t*_).

The true instantaneous reward function *R*(*s, a*) depends on the actual state and action. But for planning into the future, the agent must consider the reward as a function of its *beliefs*, which it expects to be

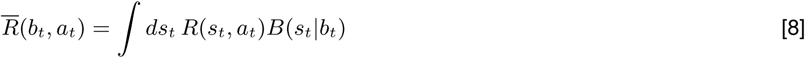

These beliefs, belief transitions 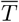, and rewards 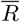 then determine the value of any policy through the recursive Bellman equation (12),

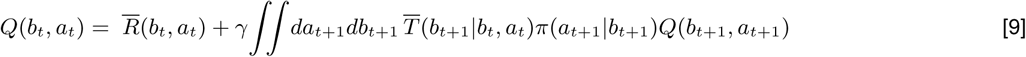

The optimal policy deterministically selects whichever action maximizes that value function given the current belief, *a*_*t*_ = argmax_*a*_*Q*(*b*_*t*_, *a*). As a generalization, here we allow actions to be sampled randomly from the softmax policy

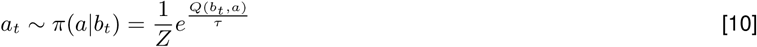

with temperature *τ* and normalization *Z*.

### Markov structure in Inverse Rational Control

The log-likelihood of the observed data ℒ (*θ*) [1] can be written as the sum of the expected complete data log-likelihood 𝒬 (*θ*) and the entropy *H* of the posterior over beliefs, ℒ (*θ*) = 𝒬 (*θ*) + *H*, as in the Expectation-Maximization (EM) algorithm (20).^‡^ Each of these terms can be decomposed into sums of transition probabilities and policies at each time, due to the Markov property. Using the graphical model structure shown in Figure 1B, we have

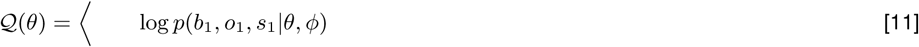

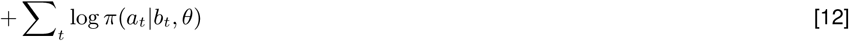

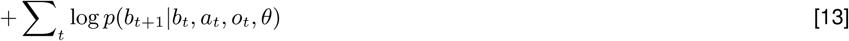

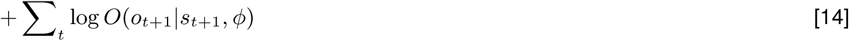

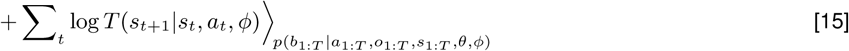

The term in [13] depends only on the parameters for the state dynamics and observations, while the policy term in [12] depends on both the dynamics and observation parameters and reward functions. The entropy *H* of the posterior can be computed similarly (see below).

Note that the true world state *s* only appears in terms with the experimental parameters *ϕ*, and does not appear with the agent’s parameters *θ* in this likelihood, because what matters to our model is not what actually happens in the world but rather what the agent *thinks* happens.

According to (69), the entropy of the posterior over beliefs can be calculated recursively as

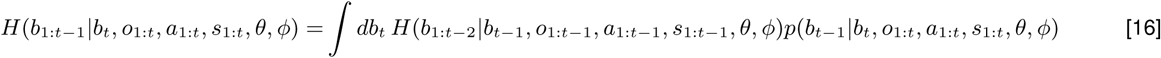

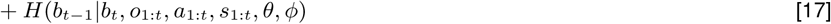

where *p*(*b*_*t*−1_ |*b*_*t*_, *o*_1:*t*_, *a*_1:*t*_, *s*_1:*t*_, *θ, ϕ*) can be calculated with Bayes rule. For the last time point, *t* = *T*, the entropy of the entire belief sequence can be obtained similarly as

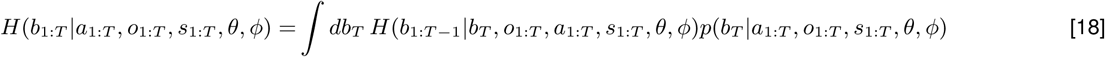

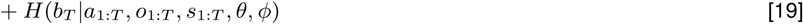

### Line search method

In small problems like the foraging task considered in the main text, we can sometimes optimize the log-likelihood function ℒ (*θ*) directly by a greedy line search method. Here we iteratively perform one-dimensional grid searches along random directions in parameter space. Once we find the optimal parameters on a line, we choose a new direction randomly from that starting point. We repeat this procedure until convergence.

### EM algorithm

The EM algorithm (20) enables us to solve for the parameters that give best explanation of the observed data, while inferring unobserved states in the model. Recall that the log-likelihood of the observed data log ℒ (*θ*) can be written as

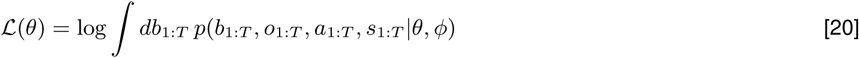

Here *θ* is a parameter vector which includes both assumptions about the world dynamics and the parameters determining the subjective magnitudes of rewards and action costs. We alternately update the parameters *θ* to improve the expected complete-data log-likelihood, and calculate the posterior over latent states based on the estimated parameters from the most recent iteration.

According to the EM algorithm, in the E-step the estimated parameters *θ*^old^ from the previous iteration determine the posterior distribution of the latent variable given the observed data *P* (*b*_1:*T*_ |*a*_1:*T*_, *o*_1:*T*_, *θ*^old^). In the M-step, the observed data log-likelihood function to be maximized reduces to

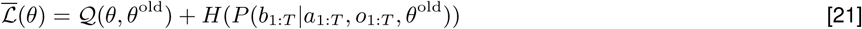

To be consistent with (70), we use 𝒬 (*θ, θ*^old^) as the auxiliary function that describes the expected complete data log likelihood, and *H*(·) is the entropy of the posterior of the latent variable. Note that *H*(·) is not a function of *θ*, and thus has a fixed value if *θ*^old^ is fixed.

The 𝒬-auxiliary function can be expressed as:

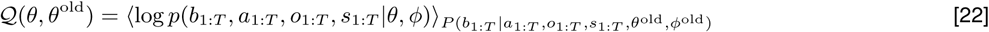

where *ϕ* are the parameters in the experimental setup that determine the world dynamics. Since *ϕ* are fixed in the experiment and known in the analysis, they do not affect the model likelihood.

The complete data likelihood *p*(*b*_1:*T*_, *a*_1:*T*_, *o*_1:*T*_, *s*_1:*T*_ |*θ, ϕ*) can be factorized into transition probabilities and policies at each time due to the Markov property. We can therefore decompose the expected complete data log likelihood 𝒬 (*θ, θ*^old^) using the graphical model structure, as described in [11–15], except now the posterior distribution over beliefs is based on the previous iteration’s parameters:

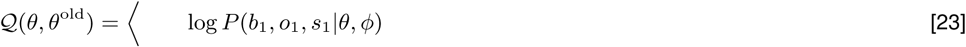

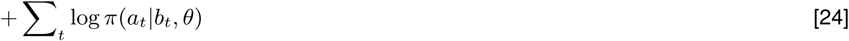

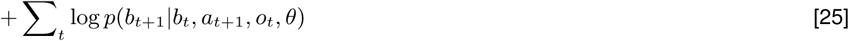

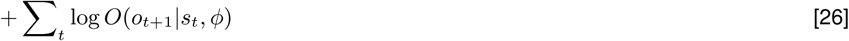

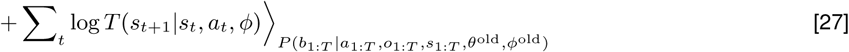

Instead of solving for the optimal *θ* in a closed form, we use gradient descent to update the parameter *θ* in the M-step.

With fixed parameters *θ*^old^ from the previous iteration, the entropy of the latent state *H*(*b*_1:*T*_ |*a*_1:*T*_, *o*_1:*T*_, *θ*^old^) is fixed. As a result, we only need to update parameter *θ* to maximize function 𝒬 (*θ, θ*^old^) in the M-step. The first term in [23] reflects the initial belief distribution, and it has a negligible contribution to when there are many time points *t*. In [25], the transition probability *p*(*b*_*t*+1_ |*b*_*t*_, *a*_*t*+1_, *o*_*t*_, *θ*) is a function of the dynamics parameters, while in [24], the policy term *p*(*a*_*t*_ |*b*_*t*_, *θ*) is a function of both the dynamic parameters and the rewards. Since the transition probability is a matrix whose elements are functions of the dynamics parameters, the gradients can be taken element-wise. We will show how the gradient of the policy function can be derived based on the *Q* value function in the next part.

Over iterations of the EM algorithm, the value of the log-likelihood ℒ (*θ*) always increases toward a (possibly local) maximum.

### Value gradient in IRC

To take gradient of the 𝒬 (*θ, θ*^old^) auxiliary function, it is critical to have the gradient of the policy function. For a softmax policy based on the value function, 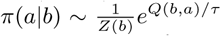, if we have the gradient of the value function with respect to the parameters, we can then obtain the gradient of the policy function using the chain rule:

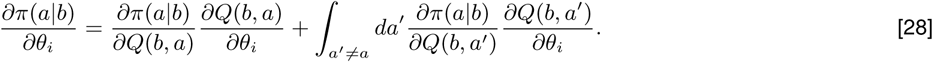

Recall that the *Q* value function for belief state-action pairs can be written as

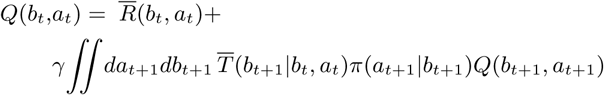

Consider now a specific element *θ*_*i*_ of the parameter vector *θ*. For a particular (*b*_*t*_, *a*_*t*_) pair, taking the derivative of both sides with respect to *θ*_*i*_, we have

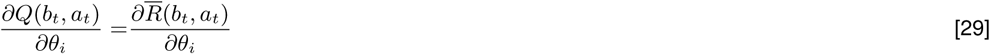

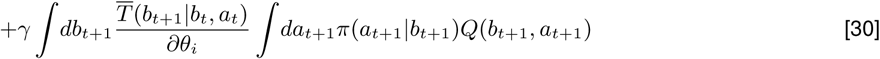

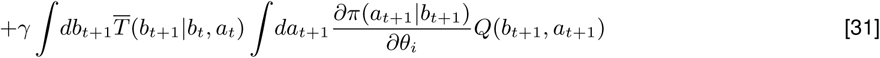

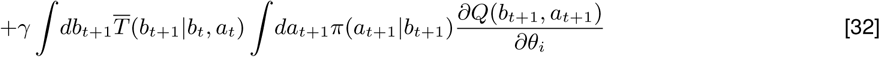

Note here 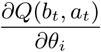 is a scalar. We define *c*_*i*_ (·) as the sum of the first two lines [29–30]:

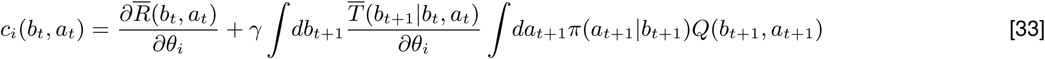

With this substitution we have

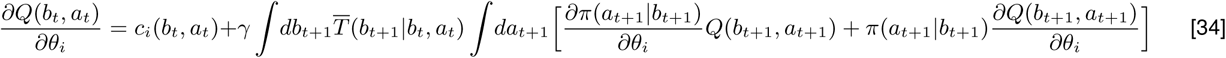

where 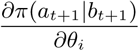 can be written as a function of 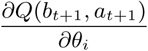 according to the chain rule [28].

Suppose there are |ℬ| distinct belief states, and |𝒜| actions. If we vectorize the matrices *Q*(*b*_*t*_, *a*_*t*_), *π*(*a*_*t*_|*b*_*t*_) and *c*_*i*_(*b*_*t*_, *a*_*t*_) over these discrete belief states and actions, denoting them as 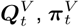 and **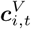** respectively, then these are vectors with length |ℬ|| 𝒜|. Equation [34] can then be rewritten as a linear function

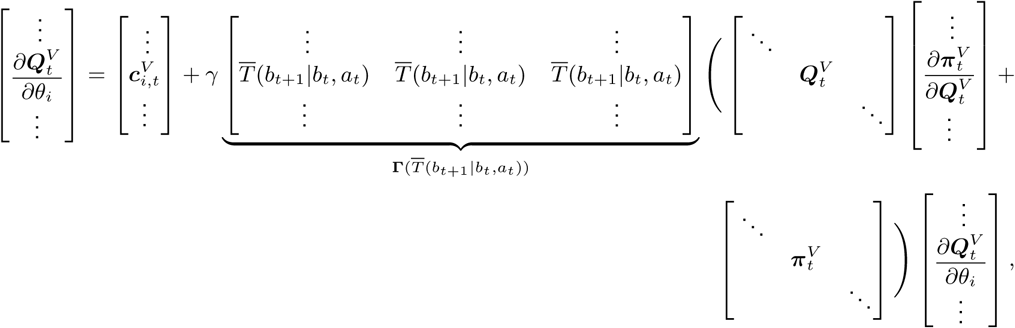

where 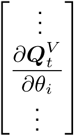 is a |ℬ|| 𝒜|× 1 vector, 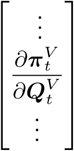 is a |ℬ|| 𝒜| × |ℬ|| 𝒜|matrix,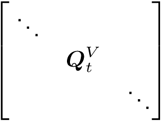 and 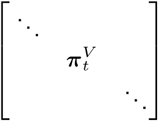 are diagonal matrices with vectors **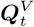** and **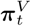** along the diagonal, and **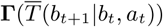** is a function of the belief transition probability 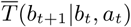. The derivative of **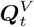** with respect to the parameter *θ*_*i*_ can then be solved as

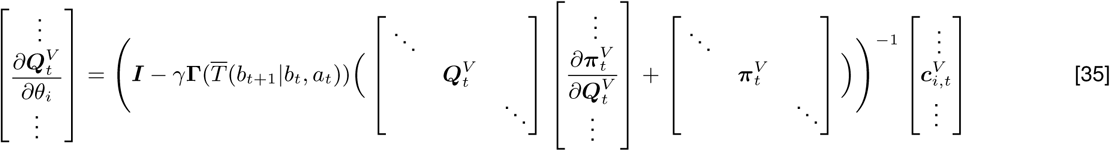

Without the brackets indicating the matrix shapes, finally we obtain

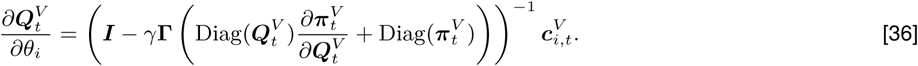

With the chain rule [28], we can obtain the gradients of the policy with respect to the parameters *θ*, which lets us calculate the gradient of the 𝒬 (*θ, θ*^old^) function in [23–27], and use them in the M-step of the EM algorithm applied to IRC. The result is an improved estimate of the agent’s internal model based on its sensory observations and actions.

### Foraging task and POMDP agent parameters

The foraging task described in the Results has two reward boxes for which the true reward availability followed a telegraph process, alternating between available and unavailable at uniform switching rates. For the two boxes, the true appearance and disappearance probabilities in one time step were 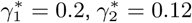 and 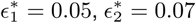.

Each box also displayed a sensory cue at each time conditioned on the reward availability, comprising five possible colors, with redder (bluer) colors indicating higher (lower) probability that food is currently available in the box. To be an interesting task, the distributions under the two states should overlap enough that the animal cannot depend primarily on the color cue to anticipate the food availability. Color values for both boxes are drawn independently at each time from a binomial distribution with five states, with mean 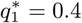 when food is available in the box, and 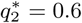 otherwise, and variance 0.96 for both of the two cases.

The target agent makes wrong assumptions about all of these parameters, acting rationally for a task where *γ*_1_ = 0.17, *γ*_2_ = 0.1, *ϵ*_1_ = 0.1, *ϵ*_2_ = 0.03, *q*_1_ = 0.45, and *q*_2_ = 0.55.

We measure gains and losses in currency of reward, *R* ≡1. In those units, our target agent incurs a subjective cost of 0.3 when pressing the button, and a cost of 0.15 when traveling. Switching between boxes requires two steps, for a total cost of 0.3. We also allow a ‘grooming’ reward *R* = 0.2 for waiting at the center location. Our agent uses a softmax policy with temperature *τ* = 0.1.

### Simulated brain

We trained a neural network to match the behavior of multiple rational agents. The target behaviors were implemented by agents that used optimal belief updates and a softmax policy with nonzero temperature. For simplicity, we discretized beliefs about reward availability for each box into *N* = 10 belief states. We defined the transition matrix in the discretized belief space by binning the continuous transition matrix 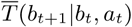. We allowed a small diffusion between neighboring bins, which reflects dynamic belief stochasticity. With the defined transition matrices and reward functions for different actions for the internal model, we can solve for the optimal softmax policy by value iteration (12).

Our neural network used one recurrently connected layer of 100 tanh units that received external inputs from the world-generated observations, agent-generated actions, and the task parameters. The recurrently connected neurons provided input to a two-layer perceptron, with 50 ReLU neurons followed by 5 policy neurons (Figure S1).

The architecture was built in PyTorch and optimized by supervised learning using gradient descent on a loss function given by the average KL-divergence between the neural network’s output policy and the teacher’s POMDP policy:

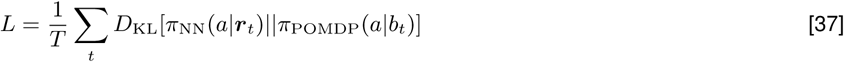

The neural network policy *π*_NN_ samples actions according to a softmax over the five output neurons 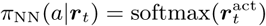. We trained the neural network to match policies with 31 teacher agents following POMDPs with different parameters. After 50 iterations of 310 batches with 1000 time points per batch, the trained neural network successfully reproduced the target beliefs within an average KL divergence of 0.0047.

**Fig. S1.**
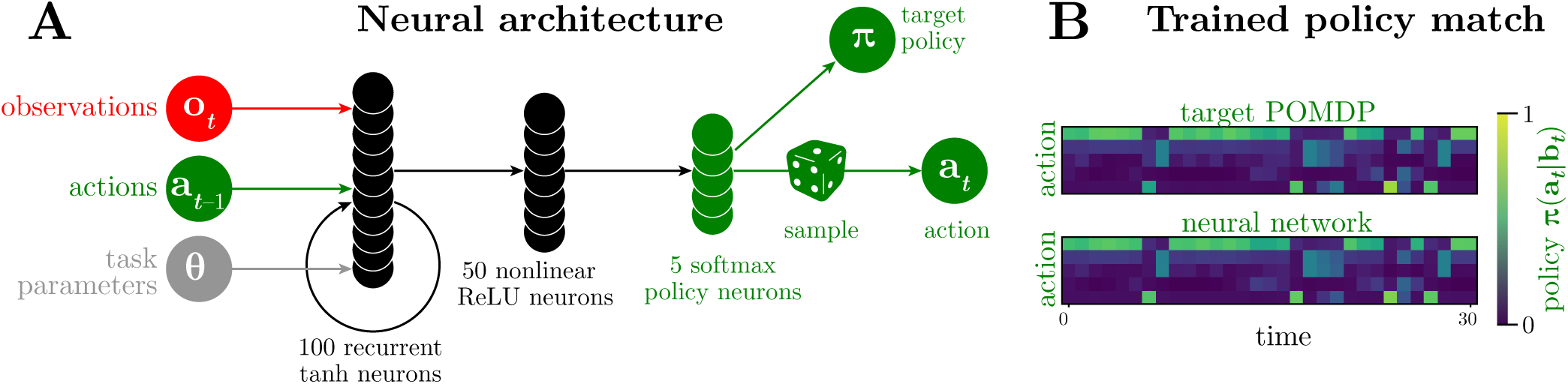
**A**: Architecture of a synthetic brain trained to behave rationally by matching the true beliefs *b* and policy *π* of a POMDP agent. The recurrent network uses 100 fully-connected neurons with a tanh nonlinearity, and the feedforward layer uses 50 ReLU neurons. There are 5 policy neurons, one for each possible action, and at each time step the network samples an action from a softmax applied to these policy neurons’ outputs. Notice that there are no hats over these quantities, because these are not estimates. **B**: Neural network reproduces the policy of a rational agent.

**Fig. S2.**
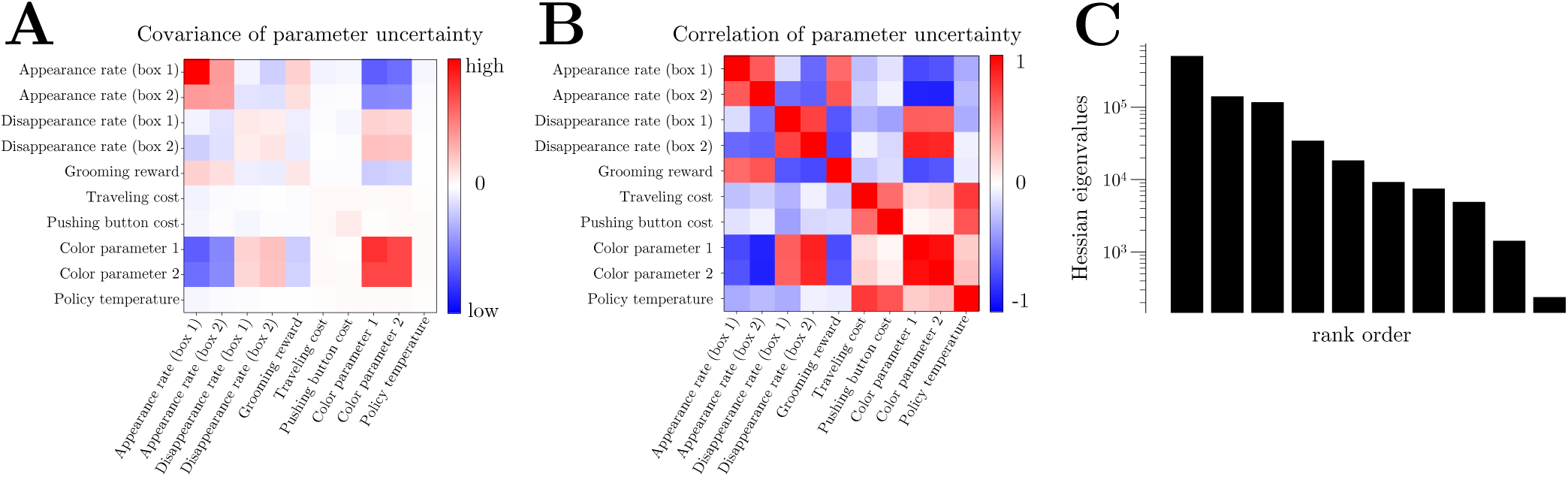
Uncertainty for the parameters fit by IRC. Confidence intervals on parameters were estimated by first using finite differences to compute the curvatures (Hessian) of the observed data log-likelihood 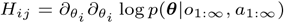, and then calculating the negative inverse ∑ = −*H*^−1^ to give the local covariance of the equivalent gaussian at the most probable value of the parameters. (**A**) The covariance matrix Σ of the local uncertainty of the model parameters. (**B**) Corresponding matrix of Pearson correlation coefficients. (**C**) Eigenvalues of the curvature matrix reveal a spectrum of uncertainties. The lowest curvature mode, *i.e.* the sloppiest direction in parameter space, has an eigenvector especially concentrated on the disappearance rate on box 2. This rate was already so low that IRC cannot tell precisely how low.

**Fig. S3.**
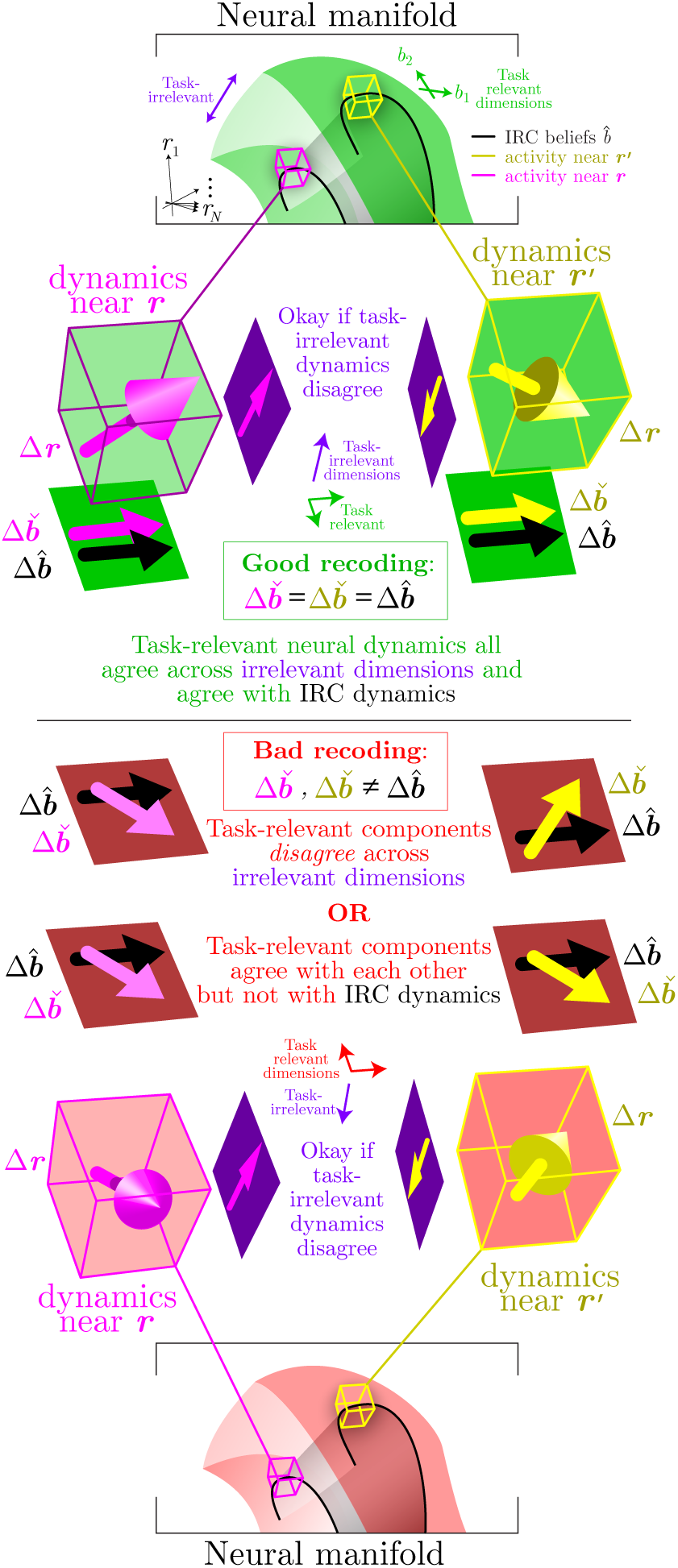
Expanding on Figure 5, the intrinsic neural dynamics define a vector field over the neural space, where the task variables evolve according to **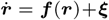** where ***ξ*** is a stochastic component that may depend on ***r***. An ideal dimensionality reduction from neural responses ***r*** (green volumes) to the task-relevant variables ***b*** must preserve the task-relevant dynamics, such that *d****ϕ***_enc_(***r***)*/dt* = ***f*** _dyn_(***ϕ***_enc_(***r***))+ ***η*** where ***η*** contains only the stochastic elements of ***r*** in the task-relevant directions. This would mean that the vector-valued updates 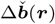 to the neural beliefs would be consistent across the (purple) task-irrelevant dimensions (yellow and magenta neural belief update vectors agree), and would also agree with the behaviorally inferred belief updates 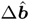 (black vectors). However, due to stochasticity, limitations in our discovery of the ideal dimensionality reduction, or a mismatch between our target behavioral model and the brain’s true model, we may find a bad representation of the task space (red volumes) for which the (yellow or magenta) task-relevant updates depend on the (purple) task-irrelevant dimensions. Two estimates of task-relevant dimensions can even have the same cross-validated encoding errors while exhibiting different dynamics with different recoding errors.

## Footnotes

* In our use of the term ‘decoding’, we are taking the brain’s perspective. The term more often reflects the scientist’s perspective, where the scientist decodes brain activity to estimate encoding quality. Instead, we reserve the term decoding to describe how neural activity affects actions: we say that the brain decodes its own activity to generate behavior.

† Estimates based on the behavioral model are consistently denoted by an up-pointing hat, 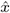, as distinguished from estimates based on the neural responses denoted by a down-pointing hat, 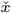, as indicated in Table S1.

‡ Unfortunately, the conventional notations in EM and reinforcement learning collide here, both using the same letter: this 𝒬 auxiliary function is denoted in the Calligraphic font to distinguish it from the state-action value function *Q* in the MDP model.

